# Assembly of photoactive Orange Carotenoid Protein from its domains unravels a carotenoid shuttle mechanism

**DOI:** 10.1101/096651

**Authors:** Marcus Moldenhauer, Nikolai N. Sluchanko, David Buhrke, Dmitry V. Zlenko, Neslihan N. Tavraz, Franz-Josef Schmitt, Peter Hildebrandt, Eugene G. Maksimov, Thomas Friedrich

## Abstract

The Orange Carotenoid Protein (OCP) is indispensable for cyanobacterial photoprotection by quenching phycobilisome fluorescence upon photoconversion from the orange OCP^O^ to the red OCP^R^ form. Cyanobacterial genomes frequently harbor, besides genes for Orange Carotenoid Proteins (OCPs), several genes encoding homologs of OCP’s N- or C-terminal domains (NTD, CTD). Unlike the well-studied NTD homologs, called Red Carotenoid Proteins (RCPs), the role of CTD homologs remains elusive. We show how OCP can be reassembled from its functional domains. Expression of *Synechocystis* OCP-CTD in carotenoid-producing *Escherichia coli* yielded violet-colored proteins, which, upon mixing with the RCP-apoprotein, produced an orange-like photoswitchable form that further photoconverted into a species spectroscopically indistinguishable from RCP, thus demonstrating a unique carotenoid shuttle mechanism. The CTD itself is a novel, dimeric carotenoid-binding protein, which effectively quenches singlet oxygen and interacts with the Fluorescence Recovery Protein, assigning physiological roles to CTD homologs and explaining the evolutionary process of OCP formation.

**One Sentence Summary:** The C-domain of cyanobacterial OCP dimerizes, binds a carotenoid, and delivers it to the N-domain forming photoactive OCP.

---

The 35 kDa photoactive Orange Carotenoid Protein (OCP) plays a central role in cyanobacterial photoprotection. Upon illumination with blue light, OCP undergoes phototransformation from the basic, orange form OCP^O^ into the red signaling form OCP^R^ (*1–4*), the latter quenching the fluorescence of phycobilisome (PBs) antennae, thus preventing excessive energy flow to the photosystems to enact non-photochemical quenching, NPQ. Besides acting in NPQ, OCP is also an efficient quencher of singlet oxygen, ^1^O_2_ (*5*). Termination of NPQ and reestablishment of full antennae capacity invokes the action of the 14 kDa Fluorescence Recovery Protein (FRP) (*1, 3, 6, 7*). However, the site where FRP binds to OCP is still controversial (*6, 8, 9*).

The X-ray crystal structure of the orange form of OCP from native cyanobacteria is known since many years (*10, 11*), whereas the red form has evaded structural characterization so far due to its lower thermodynamic stability. Very recent revolutionary progress in recombinant protein production using xanthophyll-producing *Escherichia coli* strains (*12, 13*), allowing for flexible manipulations of the OCP sequence and the nature of bound carotenoids, have greatly simplified the expression of OCP proteins for functional and structural studies (*8, 13–17*). OCP consists of two distinct structural domains, an N- and a C-terminal domain (NTD, CTD), connected via a linker that plays an important role in holding NTD and CTD together upon OCP photoactivation and domain separation (*15, 18*). A recent survey of 255 cyanobacterial genomes revealed that 87 thereof contain at least one homolog of the NTD, from which the majority (83) also contain a homolog of the CTD (*17*). This suggests that combinatorial heteromeric assemblies of isolated NTDs and CTDs could substantially expand the versatility of sensor/effector activities based on OCP’s modular architecture (*17, 19*). However, while the role of the NTD, also known as Red Carotenoid Protein (RCP) (*16*), and its homologs from the helical carotenoid protein (HCP) superfamily is well-established as phycobilisome fluorescence and singlet oxygen quenching (*19*), the role of the CTD and its multitude of homologs in numerous species is elusive.

## Structural organization of a unique carotenoid-binding protein, COCP

Upon expression of the 18 kDa OCP-CTD from *Synechocystis sp.* PCC 6803 in *E. coli* strains producing echinenone (ECN) and canthaxanthin (CAN), an unprecedented, deeply violet-colored protein was obtained (Figure 1A, inset). The protein was not photoactive, as no changes in the absorption spectrum in the visible region were observed upon strong blue-light illumination (Supplementary Figure S1A). Since it represents a novel carotenoid-binding entity distinct from other known OCP-related proteins, we named this protein C-terminal OCP-related Carotenoid Protein, COCP, to discriminate it from its N-terminal counterpart, RCP.

**Figure 1:**
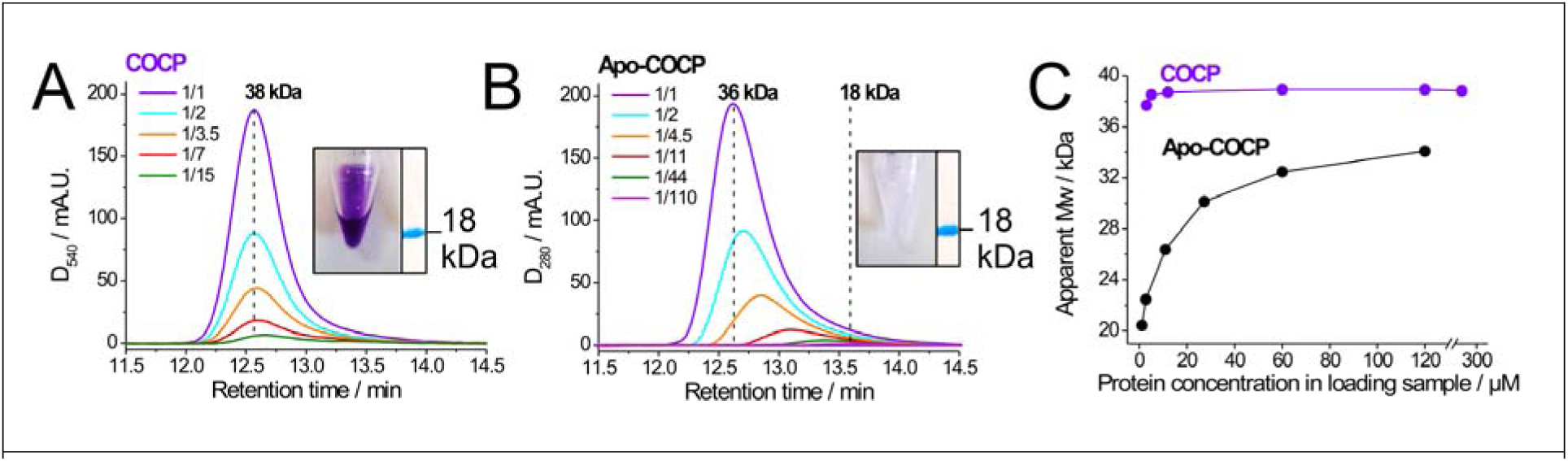
Oligomeric status of COCP and Apo-COCP determined by analytical SEC. (A) SEC elution profiles of COCP (detection: 540 nm) and (B) Apo-COCP (detection: 280 nm) at increasing dilutions as indicated (initial protein concentrations: COCP: 3.4 mg/mL; Apo-COCP: 2.1 mg/mL). Insets in (A,B) show solutions of the respective purified proteins and Coomassie brilliant blue-stained bands on SDS-electrophoregrams with molecular weight (Mw) marker indicated. (C) Dependence of the apparent Mw on protein concentration for samples from (A) and (B) estimated for COCP and Apo-COCP based on a column calibration with protein standards of known Mw. See Supplementary Methods and (8) for further details.

According to far UV-CD spectroscopy data (Supplementary Figure S2), the secondary structure of purified COCP corresponded perfectly well to the CTD folding from the latest OCP crystal structure (PDB 4XB5; residues 165-317), which contains 25 residues in α-helices (16.4 %) and 52 residues in β-strands (34.2 %). In line with this, deconvolution of the UV-CD spectrum of the COCP protein using the CDSSTR algorithm of the DichroWeb server (*20*) yielded 15 % α-helices and 32 % β-strands (NRMSD = 0.027), suggesting an unperturbed conformation. We found remarkable similarity between the secondary structure content of COCP and the unrelated α/β-folded human steroidogenic acute regulatory protein 1 (StARD1) (*21*) (Supplementary Figure S2), whose close homolog, StARD3, was found to bind the carotenoid lutein in human retina (*22*). On SDS-PAGE, the violet COCP (and the colorless apoprotein thereof, Apo-COCP) migrated as a single band of ~18 kDa (Figure 1A,B insets). However, in analytical size-exclusion chromatography (SEC), COCP and Apo-COCP eluted significantly earlier than expected for their monomers, suggesting that a higher-order oligomeric form is present (Figure 1A,B). For the violet COCP, molecular weight estimation based on column calibration resulted in a value of 38 kDa ideally corresponding to COCP dimers, with hydrodynamic radius *R_H_* = 2.80 nm (i.e., gyration radius *R_g_* = 2.17 nm for a near-spherical particle), and these dimers were stable even upon manifold sample dilution (Figure 1A,C). In contrast, we observed strong concentration-dependent dissociation in the case of Apo-COCP (Figure 1B,C), from near-dimeric (34 kDa) at higher, to almost completely monomeric forms (~20 kDa, *R_H_* = 1.94 nm) at lowest protein concentrations (Figure 1A,C). Such a dynamic behavior of Apo-COCP and the stability of the violet COCP holoprotein unequivocally indicate that the carotenoid plays a significant structural role in stabilizing the assembly of the COCP homodimer.

In striking contrast, irrespective of the presence of a carotenoid, the second known OCP module, RCP (the 19.4 kDa NTD of OCP), was monomeric with an apparent molecular mass of 20.1 kDa, almost independent of protein concentration (Supplementary Figure S3). The strong concentration dependence of Apo-COCP observed here is remarkably similar to the previously reported concentration-dependent behavior of an OCP^R^ analogue, the purple OCP^W288A^ mutant protein (*13*). This distinct property of the OCP-CTD explains the pronounced propensity of the active quenching OCP forms to self-association and may be relevant for the photoprotection mechanism.

## Spectroscopic characterization of COCP

The homodimeric arrangement, having nearly half of the carotenoid-binding cavity in each CTD, suggests that one, most likely *symmetric*, carotenoid molecule is attached in an equivalent fashion to two identical COCP monomers. Surprisingly, this was confirmed by liquid-chromatography/mass-spectrometry (LC-MS) data showing that COCP obtained from *E. coli* strains producing both ECN and CAN (Figure 2C) contains 95 % of the symmetric CAN and only 5% ECN, in strong contrast to OCP, which mainly coordinates the asymmetric ECN under identical expression conditions (Supplementary Figure S4). Therefore, most probably, each monomer within the COCP homodimer coordinates one of the two 4-keto groups of the terminal β-rings of a single CAN molecule by the specific H-bond pattern observed in the crystal structures of OCP^O^. As a result, COCP(CAN) exhibits a unique absorbance spectrum, which is broad and without vibronic substructure (Figure 2A). With a maximum at 551 nm, it is the most red-shifted from all OCP-like proteins reported to date, and even more red-shifted than the ones of RCP and of the purple OCP^W288A^ mutant described previously (*8*).

**Figure 2:**
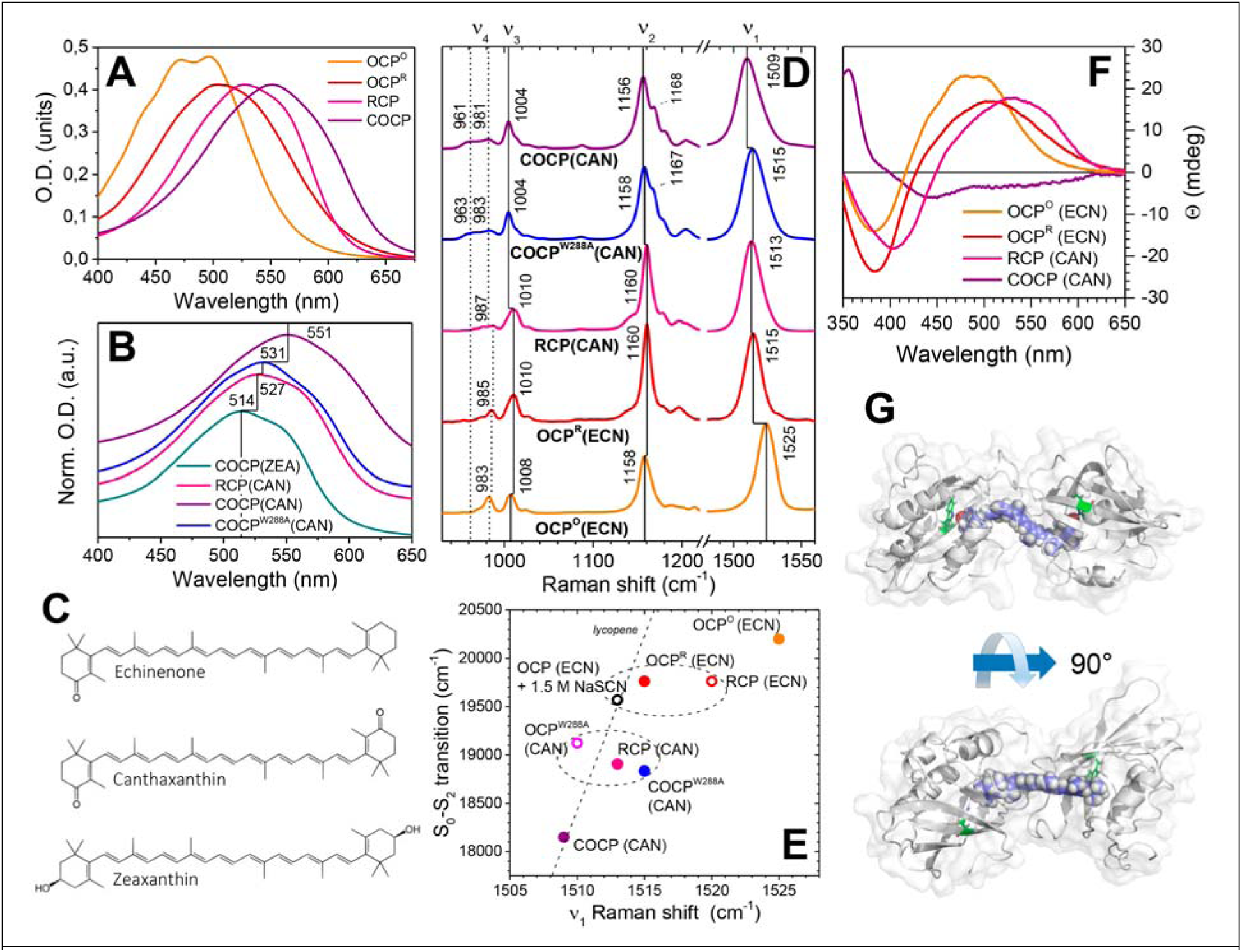
Spectroscopic properties of COCP and other OCP-related proteins. (A) Visible absorption spectra of COCP, RCP and OCP in the orange and red state (OCP^O^ and OCP^R^, respectively). For carotenoid content analysis, confer Supplementary Figure S4. (B) Absorption spectra of COCP(CAN), COCP(ZEA), COCP^W288A^(CAN) and RCP(CAN). Spectra were shifted along the y-axis for better presentation. (C) Chemical structures of echinenone (ECN), canthaxanthin (CAN) and zeaxanthin (ZEA). (D) Pre-resonance Raman spectra of the orange and red OCP forms and its carotenoid-binding domains. Numbers above the lines indicate the positions of v_1_ - v_4_ bands. (E) Correlation between the position of the S_0_→S_2_ electronic transition and the v _1_ band frequency for different OCP forms. Open circles indicate parameters taken from literature (*13, 18*). The dashed line indicates the dependency of spectral characteristics of lycopene in different solvents (*23*). (F) Vis-CD spectra of the orange and red OCP forms and its individual carotenoid-binding domains. (G) Results of a molecular dynamics simulation experiment demonstrating the structure of the COCP homodimer composed of two OCP-CTD subunits (initial coordinates from PDB 3MG1 (*11*)) coordinating one CAN molecule via H-bonds from Trp-288 in each subunit to the 4-(4’)-keto groups at the terminal β-rings. For further explanation, see text and Supplementary Materials.

Interestingly, by utilizing *E. coli* strains tailored to synthesize the symmetric carotenoid zeaxanthin (ZEA), we again obtained colored COCP protein with characteristic absorption spectra (Supplementary Figure S1), further supporting the general ability of the novel carotenoid-binding protein, COCP, to accommodate carotenoids with symmetrical β-ring structures. Of note, expression of a mutant COCP^W288A^ carrying a substitution of the critical Trp residue involved in H-bonding with the 4-keto group of the xanthophyll (*8, 13*) in a ECN/CAN-producing *E. coli* strain again yielded a CAN-coordinating protein (Supplementary Figure S4), but with a less red-shifted absorption spectrum compared to COCP, very similar to the spectrum of RCP (Figure 2B).

To get deeper insight into the chromophore structure, we scrutinized COCP by Raman (RR) and visible circular dichroism (Vis-CD) spectroscopy. RR spectra of COCP(CAN) and COCP^W288A^(CAN) were recorded and compared to the ones of other OCP variants characterized previously (*13, 18, 23*). In accordance with the established band assignment for carotenoids (*24*), the V1 band positions of COCP (1509 cm^−1^) and COCP^W288A^ (1515 cm^−1^) conform with the well-known correlation between the v_1_ band position (for OCP^O^: 1525 cm^−1^; for RCP: 1513 cm^−1^) (Figure 2D) and the spectral position of the chromophore’s S_0_-S_2_ transition inferred from the maximum of the absorption spectrum (*23, 25*), and notably places the v_1_ band COCP on a correlation line determined for the linear C_40_ carotenoid lycopene in solvents of different polarizability (*23*). Recent theoretical studies show that this correlation can be rationalized for different carotenoid-binding proteins by a combination of two effects: (i) the rotation of the β- ionone rings with respect to the conjugated system (*26*), and (ii) the local electric field in the chromophore-binding pocket (*27*). As discussed above, the preference of COCP for CAN over ECN points to the importance of H-bonding between the ketolated β-rings and Trp288/Tyr201 of COCP. Therefore, the effect of the W288A mutation (partly) disturbing this H-bonding pattern and inducing the correlated shifts between 1509 cm^−1^ and 1515 cm^−1^ band positions in RR spectra (Figure 2D) and the profound (20 nm) blue-shift in the absorbance spectra (Figure 2B), supports a structural model for COCP, in which a homodimer of two COCP subunits coordinates one CAN chromophore (Figure 2G).

Compared to OCP^O^ and RCP, the v_1_ bands of COCP and COCP^W288A^ are significantly broader, indicating a larger structural flexibility of the carotenoid in these proteins. The v_2_ bands of COCP and COCP^W288A^ are also broadened and exhibit significant splitting into two lines with another elevated shoulder on their high-energy flanges, suggesting a mixture of chromophore configurations. Significant differences can also be discriminated at the v_3_ band (composed of CH_3_-rocking vibrations) with a rather sharp peak at 1005 cm^−1^ and a smaller at 1013 cm^−1^. Also, the low energy flanges of the v_4_ band of COCP and the COCP^W288A^ mutant are elevated, a unique feature that has not been previously observed in other OCP proteins. Since the v_4_ band comprises contributions from C-H hydrogen-out-of-plane (HOOP) modes at the polyene chain, which are not or only little Raman-active if the chromophore adopts a relaxed, planar structure as in RCP (*16, 18*), the elevated Raman bands of the v_4_ region suggest a certain species with distorted or squeezed configuration of the carotenoid in COCP and COCP^W288A^, which is different from the one in OCP^O^, the latter showing the highest v_4_ band at 983 cm^−1^, but no Raman signals at the band’s low-energy flange (960 cm^−1^).

To understand the possible chirality of the carotenoid environment created by its embedment into an asymmetric protein matrix, we recorded Vis-CD spectra of COCP and other OCP variants (Figure 2F). The investigated CAN-constructs of RCP and COCP have lower ellipticities in the range of the visible absorption spectrum than all the ECN-containing variants of full-length OCP proteins, because (i) the symmetric CAN chromophore is inherently less chiral then ECN and (ii) the asymmetry of the protein environment is reduced, as suggested for the relation between OCP and RCP Vis-CD spectra (*28, 29*). In the RCP crystal structure (*16*) the chromophore is planar, with one β-ring rotated out-of-plane and the other coplanar with the polyene chain, which is most likely the reason for the low CD activity of RCP(CAN) (*28*). The low Vis-CD intensity, the red-shifted absorption, and the low-energy v_1_ mode at 1509 cm^−1^, which correlates with the spectral features of lycopene (the C_40_ carotenoid with the longest conjugated system possible), collectively indicate that in COCP, both β-ionone rings are planar with respect to the conjugated system. This also coincides with crystal structures and calculations of various red-shifted carotenoid chromophores containing planar β-rings, e.g., of crustacyanin, which endows lobster carapace with its deep bluish-black color (*27, 30*). Furthermore, the unique HOOP band pattern of COCP indicates another type of non-planarity of the conjugated system compared to the lower intensity HOOP bands of the planar chromophore in RCP, which cannot be explained by a bending of the chromophore, because this would lead to the characteristic 983 cm^−1^ band observed in OCP. Also, the deformation of the conjugated chain must be symmetric to be in accordance with the low-amplitude Vis-CD spectrum and the symmetric, homodimeric protein composition. Hence, we propose a symmetric squeezing and torsion of the conjugated chain on CAN in COCP, which is the only possible interpretation left (Figure 2G). Interestingly, the novel HOOP motif of COCP is conserved in COCP^W288A^, whereas the v_1_ band of the latter is upshifted to 1515 cm^−1^, a position similar to RCP(CAN). These spectral features indicate that the structure of the conjugated chain and the overall protein configuration is approximately the same in COCP^W288A^, but the lack of stabilizing H-bonds at the β-rings leads to a symmetrical out-of-plane relaxation of the rings. The positions of the v_1_ band and the S_0_-S_2_ transition indicate that the rotational angle of both rings must be smaller than the rotation of the β_1_-ring in RCP to bring about similar phenotypes in all spectroscopic features.

From all these spectroscopic and structural observations, we propose a structural model of COCP based on the crystal structure of the COCP coordinating CAN (PDB 3MG1, (*11*)). After optimization by molecular dynamics simulations on a rather long 600 ns timescale (see Supplementary Materials), the model (Figure 2G) represents a conformation consistent with our Vis-CD, RR, SEC, and secondary structure data, and its resultant *R_g_* value of 2.15 nm matches the one derived from SEC (2.17 nm). The model shows the CAN chromophore with coplanar arrangement of the terminal β-rings relative to the polyene chain, with the preserved H-bonding of the keto-groups with the Trp-288 residues in each COCP monomer.

## Functional properties of COCP

The FRP binding site on OCP, albeit important for understanding their interaction during termination of PBs fluorescence quenching in the context of cyanobacterial photoprotection, still remains controversial (*6, 8, 9*). To assess the possibility of a physical interaction between COCP and FRP, we employed SEC (Figure 3A) and native gel-electrophoresis (Supplementary Figure S5). As shown earlier (*8*) and above, both, FRP and COCP(CAN) eluted as 28.3 and 38 kDa dimeric proteins, respectively (Figure 3A). COCP/FRP mixtures at an excess of FRP eluted as a Peak I (Figure 3A), which is substantially shifted relative to the COCP peak and corresponds to a species with an apparent Mw of 47.3 kDa. This implies the formation of a stable COCP/FRP complex at an apparent stoichiometry, that is, however, clearly distinct from dimer/dimer (theoretically ~66 kDa). Importantly, SDS-PAGE analysis confirmed the presence of both, COCP and FRP proteins, in Peak I (Figure 3A, inset). Unfortunately, taking into account possible conformational changes upon complex formation, we cannot unambiguously delineate, whether the apparent Mw of 47.3 kDa corresponds to a stoichiometry of 2:1, 1:1, or a 1:1/2:2 mixture. Importantly, the COCP/FRP complex and its components could be separated by native gel electrophoresis (Supplementary Figure S5), which supports the conclusions from SEC data and confirms tight interaction of both, COCP and Apo-COCP, with FRP. In contrast, no evidence for such an interaction could be found for RCP/FRP mixtures analyzed by SEC. In contrast to COCP, and irrespective of the presence of carotenoid, RCP was present as a monomer at all concentrations tested (Figure 3B and Supplementary Figure S2), and no shifts in the retention time were observed at increasing concentrations of FRP (Figure 3B). Although we cannot exclude that FRP, in the context of the full-length OCP, may also contact the NTD, the CTD is individually capable of interacting tightly with FRP, suggesting that the main FRP binding site is located on the OCP-CTD.

**Figure 3:**
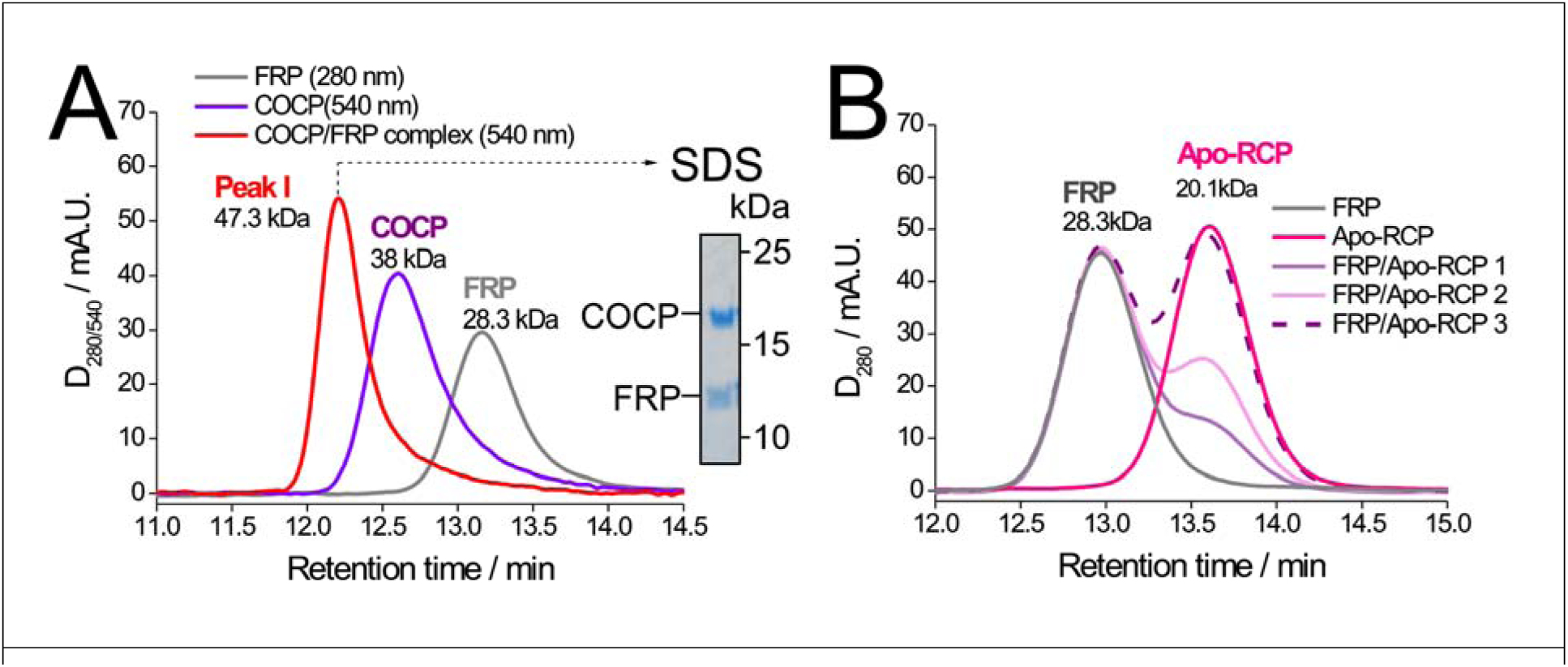
Interaction of functional modules of OCP, COCP and RCP with FRP analyzed by SEC. (A) Elution profiles of COCP (50 μM), FRP (18 μM), and a mixture of both (50 μM COCP/70 μM FRP) followed by absorbance at the indicated wavelengths. The inset shows a Coomassie brilliant blue-stained SDS-PAGE of the eluate from Peak I, with Mw markers and positions of COCP and FRP indicated. (B) Elution profile of FRP (26 μM), Apo-RCP (18 μM), or mixtures of FRP (26 μM) with increasing amounts of Apo-RCP (4.3 μM (1), 8.6 μM (2), 18 μM (3)), followed by absorbance at 280 nm. Note that profile 3 ideally coincides with the individual FRP and Apo-RCP profiles indicating no protein-protein interaction and that the presence of carotenoids in COCP and RCP does not affect their ability to interact with FRP. See Materials and methods for details.

The ability of COCP to quench ^1^O_2_ was tested by illumination of common ^1^O_2_ sensitizers, either tetra- and pentakis(cholinile) aluminium phthalocyanines (AlPCs) (31) or Rose Bengal (as used in previous ^1^O_2_ quenching assays (*5, 10*)), using Singlet Oxygen Sensor Green (SOSG) fluorescence to monitor ^1^O_2_ generation. Both, COCP and OCP caused a profound reduction of ^1^O_2_ yield (Supplementary Figure S6). Surprisingly, COCP appears to be an even better ^1^O_2_ quencher than OCP in the orange state in both assays. This ROS-quenching ability of COCP suggests a photoprotective role of OCP-CTD homologs *in vivo.*

## Reassembly of a photoactive OCP from functional modules

Since the available OCP crystal structures suggest a specific interface between NTD and CTD involving the Arg-155/Glu-244 salt bridge, an extensively H-bonded interface between the N-terminal extension (residues 3-15 of *Synechocystis* OCP) of the NTD attaching several β-sheets on the CTD (*11*), and a flexible loop of the CTD (residues 274-284) contacting the NTD from the opposite side, it seems probable that the two domains have an appreciable affinity to each other, even if not physically linked. In order to test the possibility to reassemble a photoactive OCP protein from individual structural domains, we studied the absorption spectra of mixtures of (i) Apo-COCP and RCP, and (ii) COCP and Apo-RCP by mixing constant concentrations of the carotenoid-containing forms with increasing amounts of the apoprotein forms. We did not find any changes of RCP absorption upon addition of Apo-COCP, in line with previous findings that partial proteolysis of OCP causes exclusive formation of RCP (*18*), suggesting that the carotenoid has higher affinity to the OCP-NTD (*16*). In contrast, addition of Apo-RCP to COCP caused dramatic changes of the absorption spectrum (Figure 4A) indicating rearrangement of the chromophore’s local environment. Surprisingly, the difference between the absorption spectra of COCP and mixtures with Apo-RCP is very similar to the light-minus-dark absorbance spectrum of full-length OCP (*8, 32, 33*). Indeed, the vibronic substructure characteristic for OCP in the orange form appears upon addition of Apo-RCP to COCP, revealing peaks and shoulders at 498, 468 and 437 nm (Figure 4A). However, even when Apo-RCP was present in large excess, the overall absorption changes between such a hybrid orange-like form and COCP was less than 50 % compared to the spectral changes upon OCP^O^ and OCP^R^ photoconversion. This indicates a non-homogeneous distribution of the carotenoid, in other words, the presence of red (and violet) forms in addition to an orange-like protein. The limited efficiency of the process can be at least partially explained by the dimeric nature of COCP, which implies dimer dissociation in order to interact with the NTD.

**Figure 4:**
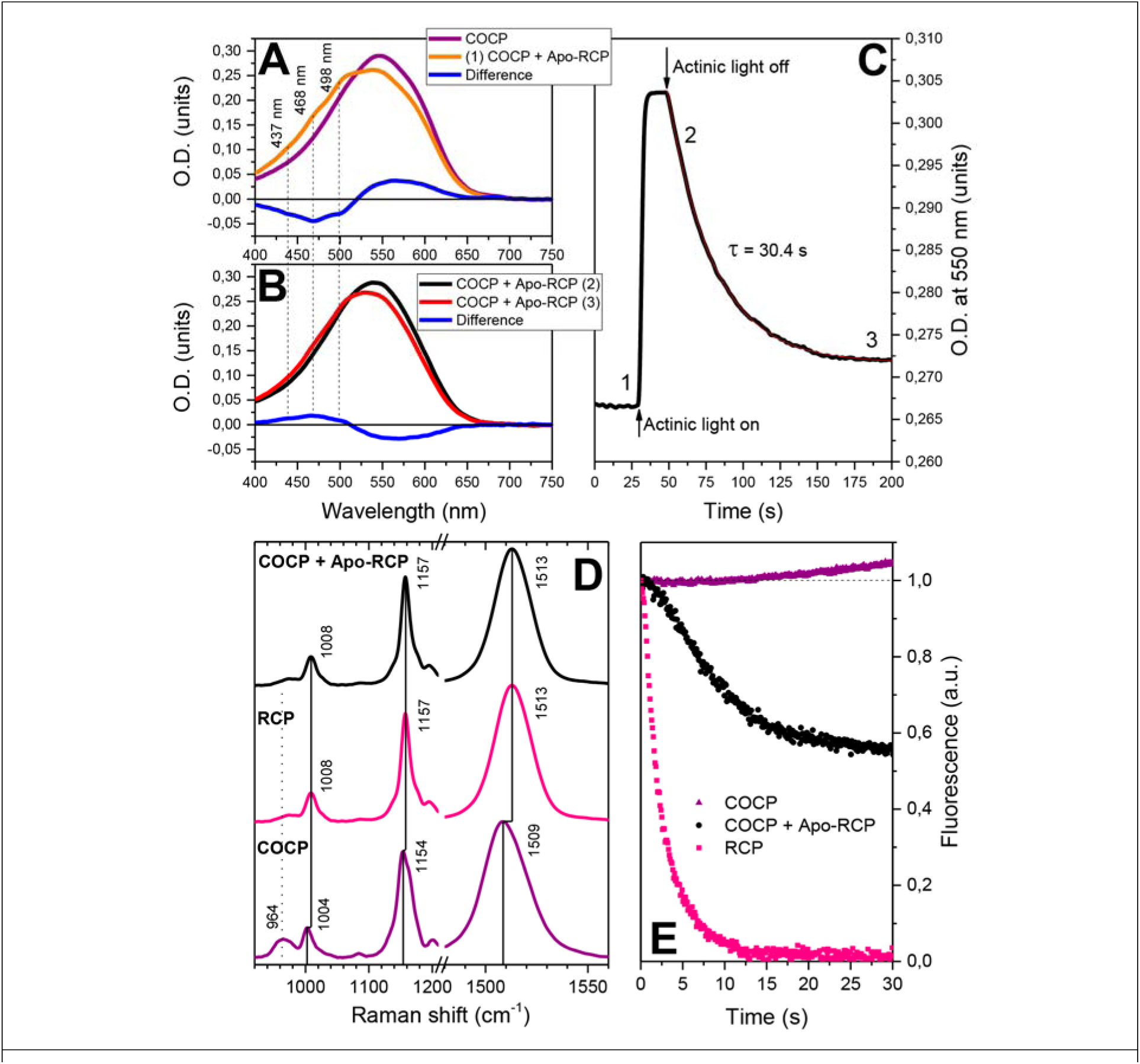
(A) Effect of Apo-RCP on COCP absorption. Apo-RCP was added until the changes of absorption were saturated. (B) Illumination of the sample in the orange-like state causes a red shift (black curve), which is partially reversible (red curve), and a part of the sample returns into the orange-like state (indicated by the shape of the difference spectra in (A) and (B)). (C) Time-course of the optical density at 550 nm upon photoconversion (actinic light: 455 nm illumination) and consequent relaxation of the sample. Numbers indicate at which moment on the time-course the numbered spectra in (A) and (B) were recorded. Absorption was measured at 25 °C and constant stirring. (D) Resonance Raman spectra of COCP, RCP and a mixture of COCP and Apo-RCP. (E) PBs fluorescence quenching *in vitro* induced by RCP (in the dark) and upon illumination of the sample containing a mixture of Apo-RCP and COCP.

The formation of an orange-like form proves that the isolated NTD and CTD of OCP interact with each other tending to form a thermodynamically stable state. Next, we tested whether this state could be affected by actinic light, which triggers conversion of OCP^O^ to OCP^R^, by measuring absorption changes at 550 nm upon illumination of the mixed sample by blue light, and obtained a kinetics typical for the absorption change of OCP^O^→OCP^R^ photoconversion and back-relaxation in the dark (Figure 4C). Importantly, in the fully converted state, the absorption of the sample was not identical to the initial (violet) COCP, but remained significantly blue-shifted. In addition, after the illumination was turned off, the absorption at 550 nm decreased monoexponentially (Figure 4C), but did not reach its initial value, indicating that parts of the sample had become trapped in the red state (Figure 4B). This phenomenon can be explained by a light-induced separation of NTD and CTD, with the carotenoid shifted into the NTD, as it is thought to occur in full-length OCP (*15*), but with the notable difference that there is no linker between the domains in our reconstructed system. Therefore, both domains might dissociate from each other upon photoconversion. As a result of this domain separation, Apo-COCP and RCP with CAN bound will be formed, which would fully explain the partial irreversibility of the spectral change observed upon such a light-triggered mechanism.

To additionally validate the hypothesis of a carotenoid shuttle mechanism between COCP and Apo-RCP, we measured RR spectra of COCP, RCP, and a mixture of Apo-RCP with COCP obtained under conditions similar to the absorption experiments shown (Figure 4A,B). Resonance Raman scattering was induced by focused 532 nm laser light, which rapidly converted any orange-like state into the red-like state. As shown in Figure 4D, the RR spectrum of the COCP/Apo-RCP mixture is identical to the one of RCP, which supports our assumption regarding the domain separation upon photoconversion and the proposed carotenoid shuttle mechanism that ultimately results in formation of RCP. Also, we tested whether the photoactive protein formed from the COCP/Apo-RCP mixture is physiologically active and studied its effect on phycobilisome fluorescence quenching. With the PBs quenching effect of RCP as a reference (Figure 4E), COCP alone did not quench PBs fluorescence. Rather, for an as yet unknown reason, a slight increase in PBs fluorescence was obtained. When kept in the dark, also the COCP/Apo-RCP mixture did not quench, but upon illumination with actinic light, a remarkable, sigmoidal decrease in PBs fluorescence was observed, substantiating our hypothesis that a quenching-competent OCP^R^-like state, and eventually RCP, is formed. Probably, the orange-like heterodimer could be stabilized by high phosphate concentration (0.8 M), which explains the lower quenching efficiency compared to RCP.

We conclude that the carotenoid, initially coordinated between two identical COCP monomers, is spontaneously inserted into the Apo-RCP module of OCP, resulting in formation of the orange-like state, which is then photoconverted into RCP. This as yet unprecedented carotenoid shuttle mechanism (Figure 5) could have profound physiological implications for OCP-CTD homologs. It seems likely that, having appeared independently in evolution, NTD and CTD precursors were combined at some point to form an ancestral protein similar to the currently known OCP. The present co-existence of OCP and its NTD and CTD homologs in cyanobacteria indicates that, at least under some circumstances, the cell requires specialized functions of these protein modules independently from each other, which additionally supports their conservation as individual entities (*17*).

**Figure 5:**
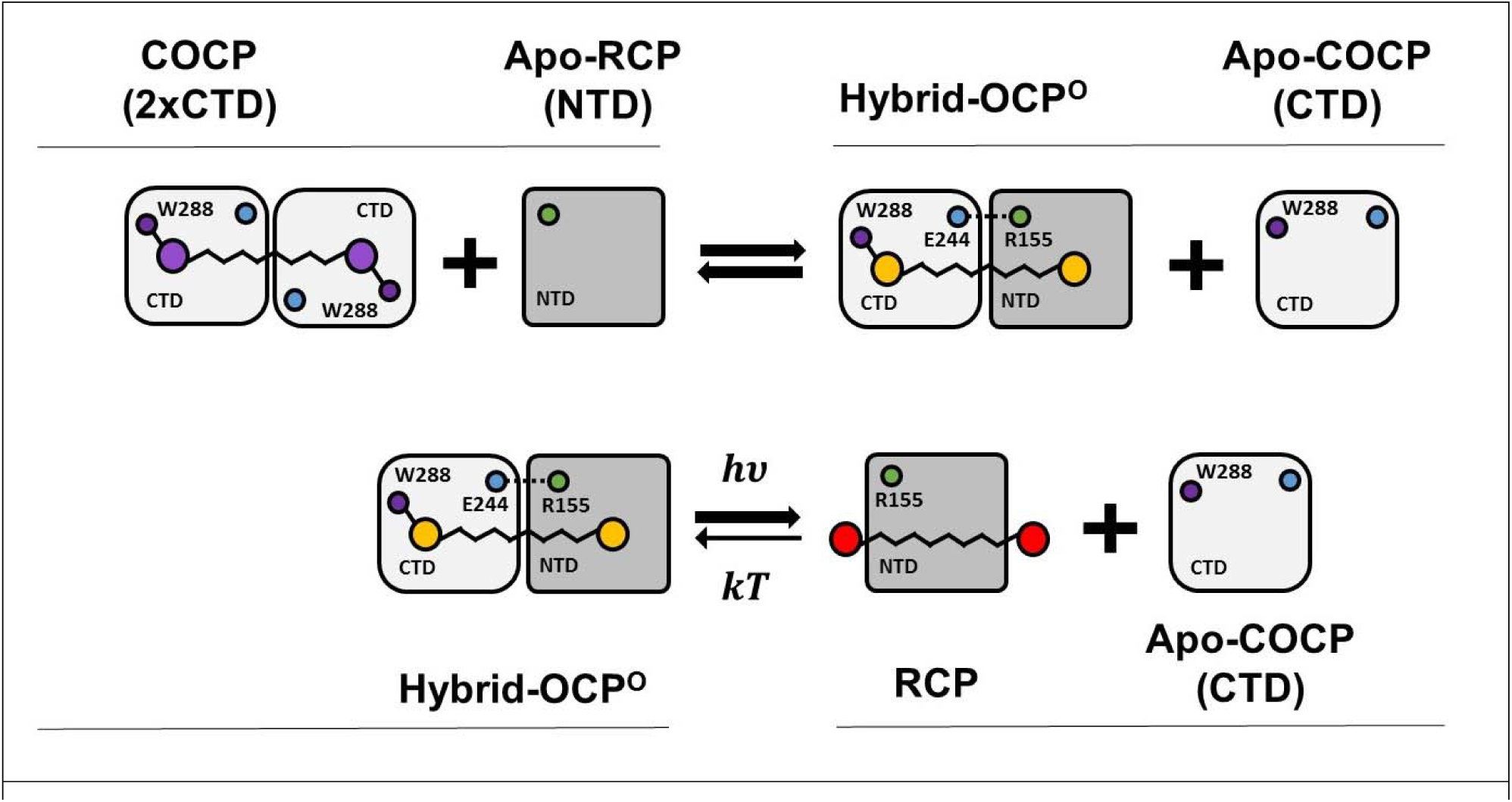
Schematic model of the interactions between COCP and the Apo-RCP and the construction of a photoactive OCP orange-like heterodimer from these proteins. COCP acts like a carotenoid carrier. Since the NTD has higher affinity to the carotenoid than the CTD, Apo-RCP accepts the carotenoid from one of the two CTD subunits of COCP, resulting in formation of an orange-like heterodimer (Hybrid OCP^O^). Upon photoactivation, the carotenoid may translocate completely into the NTD as the domain separation occurs (*15*), ultimately causing release of an Apo-COCP module and the appearance of RCP.

Carotenoids abound in photosynthetic organisms and are essential nutrients for animals. Though they partition easily into lipid compartments or membranes due to their hydrophobic nature, only very few specific carotenoid-binding proteins are known, which have the potential to maintain carotenoids in a water-soluble form and to shuttle them between membranes or cells in the organism. These rare examples include the astaxanthin-binding crustacyanin from lobster carapace (*27*), the zeaxanthin-binding GSTP1 (*34*), the lutein-binding StARD3 from human retina (*22*), and, besides OCP in cyanobacteria, the astaxanthin-binding AstaP (*35*). Thus, identification of carotenoid-coordinating proteins, which can transport their cargo from the sites of biosynthesis to deliver it at places of need, is highly relevant for understanding the physiological implications of these molecules. This study adds another important carotenoid-binding entity, OCP-CTD and its hypothetical homologous proteins of as yet orphan function (*17*), to the molecular toolkit by which organisms might purposefully employ carotenoids. Many cyanobacteria have several separate genes for CTD and NTD homologs and combinatorial heteromeric assemblies of isolated NTDs and CTDs could substantially expand the variety of sensor/effector modules based on OCP’s architecture (*17, 19*). These physiological roles may well include ROS quenching, general carotenoid transfer (e.g. from thylakoids to the outer membrane), or specific transfer to apoprotein forms of RCP homologs (or HCPs as proposed by (*19*)). In particular, the latter possibility could be a way to control the concentration of active quenchers of PBs fluorescence, and such a mechanism could be fully useful for cyanobacteria. Since COCP belongs to the NTF-2 superfamily, which also includes enzymes like ketosteroid isomerase (*36*) or naphthalene 1,2-dioxygenase (*37*), the functional spectrum could even comprise catalysis. These possibilities exponentiate the molecular toolkit made up by carotenoid-binding proteins and opens a new field of research awaiting exploration.

## Acknowledgements

Work was supported by the German Ministry of Education and Research (WTZ-RUS grant 01DJ15007 to M.M., N.N.T., F.-J.S, and T.F.), the German Research Foundation (Cluster of Excellence “Unifying Concepts in Catalysis” to T.F., D.B., and P.H) the Russian Foundation for Basic Research (grants 14-04-01536a and 15-04-01930a to V.Z.P. and E.G.M.), the Russian Science Foundation (no. 14-17-00451 to E.A.S. and E.G.M, no. 14-24-00020 to K.S.M. and D.A.L.)and the Russian Ministry of Education and Science (grant MK-5949.2015.4 to E.G.M., G.V.T. and K.E.K.). E.G.M. was supported by Dynasty Foundation Fellowship. The study was funded by RFBR and Moscow City Government (research project № 15-34-70007 «mol_a_mos»). The authors thank Prof. N. Budisa and Dr. Tobias Baumann for providing access to CD spectroscopy equipment and Dr. M. Schlangen-Ahl for technical support during LC-MS experiments.

## References

1. M. Gwizdala, A. Wilson, D. Kirilovsky, Plant Cell 23, 2631 (Jul, 2011).

2. D. Kirilovsky, C. A. Kerfeld, Biochim. Biophys. Acta 1817, 158 (Jan, 2012).

3. E. G. Maksimov et al., Photosynth. Res. 125, 167 (Aug, 2015).

4. E. G. Maksimov et al., Biochim. Biophys. Acta 1837, 1540 (Sep, 2014).

5. A. Sedoud et al., Plant Cell 26, 1781 (Apr 18, 2014).

6. C. Boulay, A. Wilson, S. D’Haene, D. Kirilovsky, Proc. Natl. Acad. Sci. U. S. A. 107, 11620 (Jun 22, 2010).

7. M. Gwizdala, A. Wilson, A. Omairi-Nasser, D. Kirilovsky, Biochim. Biophys. Acta 1827, 348 (Mar, 2013).

8. N. N. Sluchanko et al., Biochim. Biophys. Acta 1858, 1 (2017).

9. M. Sutter et al., Proc. Natl. Acad. Sci. U. S. A. 110, 10022 (Jun 11, 2013).

10. C. A. Kerfeld et al., Structure 11, 55 (Jan, 2003).

11. A. Wilson et al., J. Biol. Chem. 285, 18364 (Jun 11, 2010).

12. C. B. de Carbon, A. Thurotte, A. Wilson, F. Perreau, D. Kirilovsky, Science Rep. 5, 9085 (2015).

13. E. G. Maksimov et al., Photosynth. Res. 130, 389 (Dec, 2016).

14. A. Thurotte et al., Plant Physiol. 169, 737 (Sep, 2015).

15. S. Gupta et al., Proc. Natl. Acad. Sci U. S. A. 112, E5567 (Oct 13, 2015).

16. R. L. Leverenz et al., Science 348, 1463 (Jun 26, 2015).

17. M. R. Melnicki et al., Mol Plant 9, 1379 (Oct 10, 2016).

18. R. L. Leverenz et al., Plant Cell 26, 426 (Jan, 2014).

19. R. López-Igual et al., Plant Physiol. 171, 1852 (Jul, 2016).

20. L. Whitmore, B. A. Wallace, Nucl. Ac. Res. 32, W668 (2004).

21. N. N. Sluchanko, K. V. Tugaeva, Y. V. Faletrov, D. I. Levitsky, Prot. Expr. Purif. 119, 27 (2016).

22. B. Li, P. Vachali, J. M. Frederick, P. S. Bernstein, Biochemistry 50, 2541 (2011).

23. E. Kish, M. M. Pinto, D. Kirilovsky, R. Spezia, B. Robert, Biochim. Biophys. Acta 1847, 1044 (Oct, 2015).

24. S. Schlücker, A. Szeghalmi, M. Schmitt, J. Popp, W. Kiefer, J. Raman Spectr. 34, 413 (2003).

25. M. M. Mendes-Pinto et al., J. Phys. Chem. B 117, 11015 (2013).

26. Y. Mori, Chem. Phys. Letters 652, 184 (2016).

27. A. P. Gamiz-Hernandez, I. N. Angelova, R. Send, D. Sundholm, V. R. I. Kaila, Angew. Chem. Intl. Ed. 54, 11564 (2015).

28. P. Chabera, M. Durchan, P. M. Shih, C. A. Kerfeld, T. Polivka, Biochim. Biophys. Acta 1807, 30 (Jan, 2011).

29. J. D. King, H. Liu, G. He, G. S. Orf, R. E. Blankenship, FEBS Lett. 588, 4561 (Dec 20, 2014).

30. M. Cianci et al., Proc. Natl. Acad. Sci U. S. A. 99, 9795–9800 (2002).

31. E. G. Maksimov, D. A. Gvozdev, M. G. Strakhovskaya, V. Z. Paschenko, Biochemistry (Moscow) 80, 323 (2015).

32. E. G. Maksimov et al., Biophys. J. 109, 595 (Aug 4, 2015).

33. A. Wilson et al., Proc. Natl. Acad. Sci. U.S.A. 105, 12075 (August 19, 2008, 2008).

34. P. Bhosale et al., Journal of Biological Chemistry 279, 49447 (November 19, 2004, 2004).

35. S. Kawasaki, K. Mizuguchi, M. Sato, T. Kono, H. Shimizu, Plant Cell Physiol. 54, 1027 (July 1, 2013, 2013).

36. H.-S. Cho, G. Choi, K. Y. Choi, B.-H. Oh, Biochemistry 37, 8325 (1998).

37. B. Kauppi et al., Structure 6, 571 (1998).

38. N. Misawa et al., J. Bacteriol. 177, 6575 (Nov, 1995).

39. S.-K. Choi et al., Marine Biotechnol. 7, 515 (2005).

40. G. Britton, in Carotenoids G. Britton, S. Liaaen-Jensen, H. Pfander, Eds. (Birkhäuser, Basel, CH, 1995), vol. 1B, pp. 376.

41. F. Velazquez Escobar et al., Biochemistry 53, 20 (Jan 14, 2013).

42. M. J. Abraham et al., SoftwareX 1-2, 19 (2015).

43. B. Fernández-González, G. Sandmann, A. Vioque, J. Biol. Chem. 272, 9728 (April 11, 1997, 1997).

